# Computational Profiling and Pathogenicity Assessment of VHL Missense Mutations

**DOI:** 10.64898/2026.10.17.759794

**Authors:** Ruchir Rakesh, Edene Levine, Wilma Roch, Ana Ciochina

## Abstract

Mutations in the VHL tumor suppressor gene promote tumorigenesis by stabilizing HIF-1*α*. In this study, we investigated the functional impact of VHL missense mutations using AlphaMissense pathogenicity scores, protein stability simulations, and molecular dynamics. AlphaMissense identified high pathogenicity scores for residues 60-200, including the *β*-domain and HIF-binding domain, suggesting these regions are critical for VHL’s tumor suppressor function.

We categorized the mutations into two groups: those that primarily destabilize protein folding and those that likely disrupt VHL-HIF-1*α* binding. FoldX stability simulations revealed high ΔΔG values for both groups, with proline substitutions exerting significant structural stress. Structural modeling confirmed that these mutations disrupt secondary structures, particularly *β*-sheets.

Molecular dynamics simulations showed that the VHL monomer is flexible and unstable in several regions, whereas the VHL-HIF complex, particularly the HIF-binding domain, exhibits enhanced stability. Mutational scanning revealed that bulky amino acid substitutions, such as tryptophan, cause notable destabilization.

Interestingly, some mutations predicted to be structurally stable were still classified as pathogenic by AlphaMissense and validated in the literature, suggesting that pathogenicity may arise from subtle functional impairments rather than just structural instability.

These findings underscore the value of combining machine learning, structural simulations, and experimental data to predict and interpret the pathogenicity of VHL mutations.

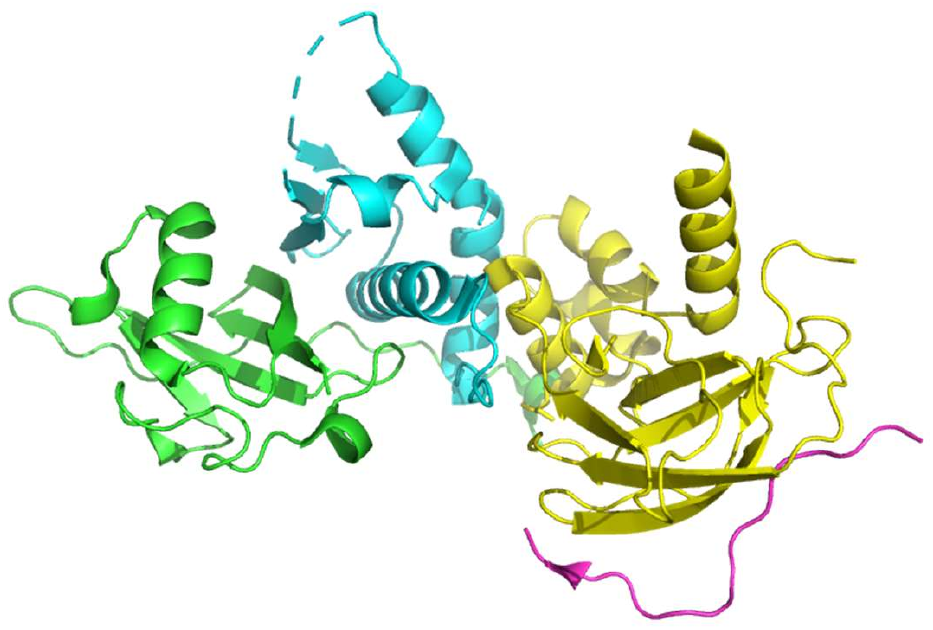

## 1 Introduction

Von Hippel-Lindau disease (VHL) is a hereditary cancer syndrome caused by mutations in the VHL tumor suppressor gene. VHL plays a crucial role in cellular responses to hypoxia. VHL is part of an E3 ubiquitin ligase complex that includes Elongin B, Elongin C, Cullin 2 and RBX1. Under normoxic conditions, the VHL complex binds to hydroxylated hypoxia-inducible factor-1*α* (HIF-1*α*), whereupon HIF-1*α* is ubiquitinated and subsequently proteasomally degraded. Under hypoxic conditions, such as those commonly found in tumors, HIF-1*α* is no longer hydroxylated by prolyl hydroxylase domain enzymes (PHDs) and therefore cannot bind to VHL. As a result, HIF-1*α* escapes degradation, accumulates in the cell, translocates to the nucleus, and functions as a transcription factor to activate genes involved in processes such as angiogenesis, glycolysis, and erythropoiesis. These pathways are frequently upregulated in cancer and promote tumor growth and metastasis. Thus, the interaction between VHL and HIF-1*α* is essential for the maintenance of cellular homeostasis (Figure 1). Importantly, this mechanism can also be triggered independently of oxygen levels if VHL is non-functional, for instance due to a pathogenic mutation [1].

**Figure 1:**
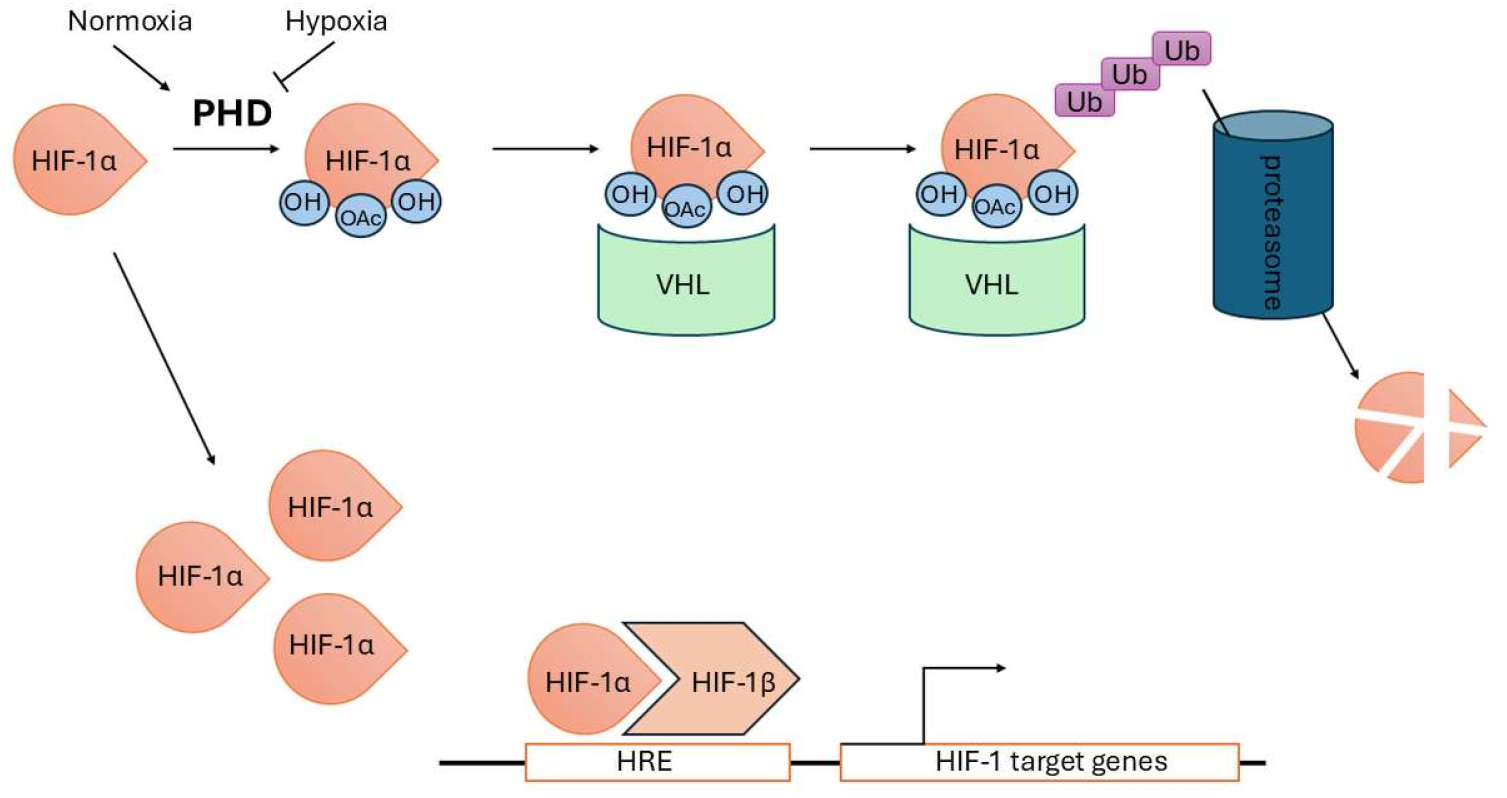
Under normoxic conditions, prolyl hydroxylases (PHDs) hydroxylate HIF-1*α*, facilitating its binding to the VHL complex. This interaction leads to ubiquitination and proteasomal degradation of HIF-1*α*. In hypoxic conditions, the activity of PHDs is inhibited due to reduced oxygen availability, preventing the hydroxylation of HIF-1*α*. As a result, HIF-1*α* accumulates in the cell, translocates to the nucleus, and activates the transcription of target genes involved in processes such as angiogenesis, glycolysis, and erythropoiesis, functioning as a transcription factor. A mutation in pVHL leads to the same effect, allowing HIF-1*α* to accumulate even under normoxic conditions.

The *β*-domain of VHL binds to the hydroxylated proline residues in the HIF-1*α* region between amino acids 532 and 585. VHL loss-of-function mutations are a common mechanism in cancers such as renal cell carcinoma (RCC), cerebellar hemangioblastoma (CHB), retinal angioma (RA), and pheochromocytoma (PCC) [2].

In this study, computational methods are used to analyse VHL and its mutations, focusing on how specific genetic alterations affect protein folding, stability, and interaction with HIF-1*α*. To assess how specific VHL mutations impact protein structure and function, we used FoldX for stability prediction, MutateX for mutational analysis, and CABSFlex for protein dynamics simulations. The aim of these analyses is to gain a deeper understanding of how mutational changes disrupt the VHL-HIF interaction. This could improve the prediction of pathogenicity and provide further insights into the molecular mechanisms of VHL-related tumorigenesis. The findings could contribute to more effective diagnostic and therapeutic strategies for VHL-related cancers.

## 2 Methods

### 2.1 Structure Selection and Variation

For the computational investigation of the VHL protein, accurate protein structure selection is essential. Experimental and in silico model structures are available from the Protein Data Bank (PDB) and the AlphaFold Database. To identify the most suitable structure for further applications PDBminer was used, an open-source software package [3]. The UniProt ID P40337 was used as input for the PDBminer algorithm, which generated a quality-ranked output table to facilitate the selection of the most appropriate VHL structure.

The output consisted of one AlphaFold-predicted structure, seven electron microscopy (EM) structures, and 108 X-ray crystallography-solved structures. The structures were first filtered by resolution, applying a cutoff of 2.5 Å. This step narrowed the selection to 61 structures. Subsequently, to focus on the VHL-HIF interaction, structures containing HIF as a chain were prioritized. This resulted in six relevant structures: 4AJY, 6GMR, 6GFX, 1LM8, and 1LQB.

Each of these structures was then evaluated to ensure that all relevant residues (typically spanning residues 60-200) were present and intact. No missing residues were found in the core protein region of any of these six structures. The final selection was based on the highest resolution, lowest R-value Free, and confirmation that the structure represented the wild-type protein. Among the six structures, 4AJY was chosen due to its resolution of 1.74 Å, an R-value Observed of 0.192, and an R-value Free of 0.232.

For variant selection, experimentally identified missense mutations from the literature [1, 4, 5] were compiled and cross-referenced with disease associations and variant information available in UniProt. Additionally, AlphaMissense was used to predict the pathogenicity of the selected mutations. The mutations were chosen based on their direct involvement in the VHL-HIF interaction and their potential impact on protein folding and stability. Only mutations previously reported in the literature, documented in UniProt, and predicted to be likely pathogenic by AlphaMissense were included in the study.

The selected mutations were as follows: Mutations within the protein that may affect its folding or stability include **N78S, Y98H, F119L, F136L, R167Q**, and **I151T**. Additionally, mutations that destabilize the interaction with HIF were identified, such as **L89P, L118P, N131T**, and **Q132P**. Furthermore, mutations that directly impact the HIF interaction at the binding site are **W88R, S111N**, and **H115P** (Figure 2).

**Figure 2:**
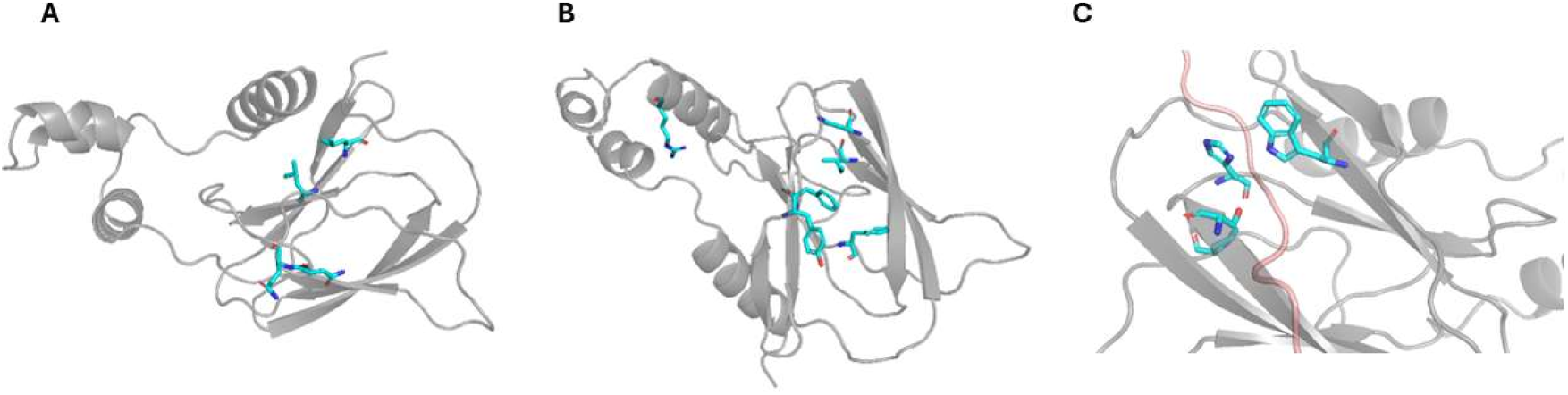
VHL is shown in grey and HIF-1*α* in red. Missense mutations within VHL, identified in the literature, are highlighted in blue. A, mutated residues that could destabilize binding to HIF-1*α*. B, mutated residues within the protein that may affect folding and stability. C, mutated residues directly impacting binding to HIF-1*α*.

The 4AJY structure comprises four chains: VHL, Elongin B, Elongin C, and HIF. For the purposes of this study, Elongin B and Elongin C were removed using PyMOL. The VHL monomer and its complex with HIF were then prepared as separate PDB files for further analysis.

### 2.2 Methods Of Stability and Simulation of the VHL Monomer

#### 2.2.1 Mutation Classification and Grouping for the Monomer

Missense mutations in the VHL monomer were divided into three groups based on their likely functional impact. Batch 1 includes mutations that are predicted to affect protein folding and structural stability - N78S, Y98H, F119L, F136L, I151T, R167Q. Batch 2 consists of mutations that are likely to disrupt the interaction between pVHL and HIF-1*α* - L89P, L118P, N131T, Q132P. This classification was informed by published experimental evidence and known disease associations [6, 7].

#### 2.2.2 Stability Simulation Using FoldX for the Monomer

To evaluate the effect of mutations on the stability of the VHL monomer on Batch 1 and 2, the FoldX software suite was employed. The wild-type crystal structure of VHL (PDB ID: 4AJY) served as the template. Each mutation was introduced using the BuildModel function in FoldX. Prior to introducing mutations, energy minimization was applied to relax the wild-type structure. After mutating, the same minimization protocol was used to ensure consistent comparison. FoldX calculated the change in folding free energy (ΔΔG), with positive values indicating destabilization and negative values suggesting stabilization.

#### 2.2.3 Structural Visualization Using PyMOL for the Monomer

To complement the energy-based predictions, PyMOL was used for structural modeling and visualization of wild-type and mutant VHL structures. This allowed visual inspection of residue environments, hydrogen bonding networks, hydrophobic packing, and steric clashes. Focus was given to proline substitutions and their effects on *β*-sheet integrity and local geometry. Distances between atoms were measured to evaluate packing disruptions, especially those under 4 Å.

#### 2.2.4 Amino Acid Property Analysis For Monomer

Wild-type and mutant amino acids were compared using a standard chart of physicochemical properties, including hydrophobicity, polarity, and structural flexibility. This comparison helped interpret the ΔΔG results and identify patterns in mutation-induced instability. The data source was Technology Networks’ essential amino acids chart [8].

### 2.3 Methods Of Stability and Simulation of the VHL-HIF1*α* Complex

#### 2.3.1 Mutation Classification and Grouping

Missense mutations in the VHL complex were divided into two groups based on their likely functional impact and molecular positioning. Batch 1 was only studied for the monomer as it was expected to affect only folding not binding. Batch 2 consists of mutations that are likely to disrupt the interaction between pVHL and HIF-1*α* - L89P, L118P, N131T, Q132P but aren’t close to the interface. Batch 3 mutations - W88R, S111N, and H115P - affect pVHL-HIF binding directly by being close to the interface. This classification was informed by published experimental evidence and known disease associations [6, 7].

#### 2.3.2 Stability Simulation Using FoldX

To study how mutations affect the stability of the VHL complex in Batches 2 and 3, the FoldX software suite was used. First, the crystal structure of the wild-type VHL protein (PDB ID: 4AJY), keeping only the V (chain A) and H (chain D) components, while removing chains B and C and all solvent molecules is used.

Mutations were then introduced one at a time using the *BuildModel* command in FoldX. This included the -I flag to specify interface residues and the -B flag to calculate changes in binding energy between the V and H chains. After each mutation, the same energy minimization process was repeated to ensure fair comparisons between wild-type and mutant structures.

FoldX calculated the change in folding free energy (ΔΔG) for each mutation. A positive ΔΔG value indicates that the mutation likely destabilizes the complex, while a negative value suggests increased stability.

#### 2.3.3 Structural Visualization Using PyMOL

To enhance the energy-based predictions, PyMOL was employed for structural modeling and visualization of both wild-type and mutant VHL structures. This approach facilitated the direct examination of residue environments, hydrogen bonding networks, hydrophobic interactions, and potential steric clashes. Particular attention was given to the impact of proline substitutions on *β*-sheet stability and local geometry. Distances between atoms were measured to assess packing disruptions, especially those within 4 Å.

#### 2.3.4 Amino Acid Property Analysis for Monomers

A comparative analysis of wild-type and mutant amino acids was conducted using a standard chart of physicochemical properties, which included hydrophobicity, polarity, and structural flexibility. This comparison provided insights into the ΔΔG results and helped identify trends in mutation-induced instability. The data for this analysis were sourced from Technology Networks’ essential amino acids chart [8].

### 2.4 Molecular Dynamics of the VHL Monomer and VHL-HIF Complex and Ensemble Mutational Investigation on the Monomer

#### 2.4.1 Generation of an Ensemble of Conformations for the Wild-Type Structure of the VHL and VHL-HIF Complex

The simulations were performed using CABSFlex [9] in the pupil terminal. The protocol involved parameters that extracted 10 medoids, conducted 10 Monte Carlo cycles, and included 10 annealing cycles. A verbosity level of 3 was selected to balance computational detail and runtime, and a random seed value of 10 was set to ensure reproducibility. All atoms were included in the simulations. This methodology was applied to both the VHL monomer and the VHL-HIF complex.

The Monte Carlo method, which iteratively samples conformations, is well-suited for predicting protein structural dynamics. The inclusion of 10 annealing cycles allowed for a more thorough exploration of the conformational landscape at different temperatures, increasing the likelihood of identifying stable structural states. A verbosity level of 3 provided sufficient detail on intermediate conformations and energy scores without compromising computational efficiency. The fixed seed ensured that the simulations could be replicated consistently.

#### 2.4.2 Structural Visualization of Dynamics Using PyMOL

For each system (monomer and complex), the program generated three output folders, one of which contained the PDB files of the 10 resulting structural models. These models were visualized in PyMOL [10] to illustrate the dynamic behavior of the protein. All models were aligned to the monomer structure for comparative analysis. In addition, the simulation output included RMSF (Root Mean Square Fluctuation) plots. RMSF quantifies atomic displacement from the mean position over time, making it a valuable metric for distinguishing between flexible and structurally stable regions within the protein.

#### 2.4.3 Mutational Self-Scan Using the CABSFLEX Model

A self-scan was performed using MutateX [11] and FoldX [12] to evaluate the local quality of the models and determine if they are suitable for assessing mutations at different sites within the protein. To conduct the self-scan on model 8, the -s option in MutateX was used, which eliminated the need for a position list (poslist). This allowed the program to scan the entire protein structure. The results from the self-scan were aggregated into a file containing the ΔΔG energy values for each residue.

#### 2.4.4 Ensemble mutational scan

MutateX [11] and FoldX [12] were applied to analyze all the CABSFlex models as an ensemble. For this analysis, a position list (poslist) of all mutation sites was provided. The output, located in a dedicated folder, included the ΔΔG estimates for the mutations across each model. These results were then extracted into a CSV file and visualized as a heatmap to represent the variation in mutation stability across the ensemble.

## 3 Results

### 3.1 Literature-Based Selection of Mutations Supported by AlphaMissense Predictions

VHL gene alterations previously described in the literature and originally identified in cancer patients with VHL disease were selected for analysis. These mutations are located in regions of the protein that may affect its folding or stability (N78S, Y98H, F119L, F136L, R167Q and I151T). Additionally, some mutations destabilize the interaction with HIF, either directly or indirectly (L89P, L118P, N131T, Q132P, W88R, S111N and H115P). The predicted pathogenicity of these mutations was assessed using AlphaMissense. All these mutations have AlphaMissense scores greater than 0.87, indicating a high likelihood of being pathogenic (Table 1).

**Table 1:** Mutations identified from literature and classified as likely pathogenic by AlphaMis-sense.

| Mutation | AlphaMissense Score | Prediction |
| --- | --- | --- |
| N78S | 0.8815 | Likely Pathogenic |
| Y98H | 0.9734 | Likely Pathogenic |
| F119L | 0.9988 | Likely Pathogenic |
| F136L | 0.9969 | Likely Pathogenic |
| R167Q | 0.8791 | Likely Pathogenic |
| I151T | 0.9960 | Likely Pathogenic |
| L89P | 0.9939 | Likely Pathogenic |
| L118P | 0.9919 | Likely Pathogenic |
| N131T | 0.9632 | Likely Pathogenic |
| Q132P | 0.9439 | Likely Pathogenic |
| W88R | 0.9990 | Likely Pathogenic |
| S111N | 0.9764 | Likely Pathogenic |
| H115P | 0.9807 | Likely Pathogenic |

Within the structural core of the VHL protein lies the so-called “aromatic tetrahedron,” a conserved motif that stabilizes the *β*-domain through hydrophobic and aromatic interactions. Disruption of this structural element compromises both the stability of the protein and its ability to bind HIF. According to the literature, residues F119 and F136 play a central functional role within this motif [1]. AlphaMissense values of 0.99 emphasize the importance of these residues in the aromatic tetrahedron for the stability and function of VHL.

Mutations at the VHL–HIF-1*α* interface (W88R, S111N, and H115P) also show high AlphaMis-sense pathogenicity scores (*>*0.97), suggesting potential functional disruption. Interestingly, although Y98 is not part of the core binding interface, it is crucial for interacting with hydroxylated proline residues on HIF-1*α*, a key step in substrate recognition and subsequent ubiquitination and degradation by the VCB E3 ligase complex.

In addition, variants such as L89P, L118P, N131T, and Q132P appear to impair HIF-1*α* binding indirectly (pathogenicity score *>*0.94). Despite not being located directly at the interface, these mutations likely alter local folding or structural dynamics near the interaction site.

Among these, L89P, L118P, N131T, Q132P, F136S, and I151T have been reported as most destabilizing mutations within the first domain in the literature [1]. These variants also exhibit high pathogenicity scores, particularly L89P, L118P, and F136S, each scoring 0.99, further supporting their functional relevance.

It is noticeable that the AlphaMissense pathogenicity score is generally very high at values between residues 60 and 200 of the VHL protein. This is where the *β*-domain and the HIF-binding domain are located, which are responsible for the tumor suppressor function of VHL. The accumulation of likely pathogenic variants within this segment further demonstrates its functional importance and structural susceptibility.

### 3.2 Simulated Results on the Stability of the VHL Monomer

To evaluate the impact of each mutation on the stability of the VHL monomer, ΔΔG values were calculated using FoldX. These energy changes reflect the destabilizing potential of each variant, with higher ΔΔG values indicating greater structural disruption. The results are summarized in Table 2, which presents two mutation groups categorized by their predicted functional consequences.

**Table 2:**
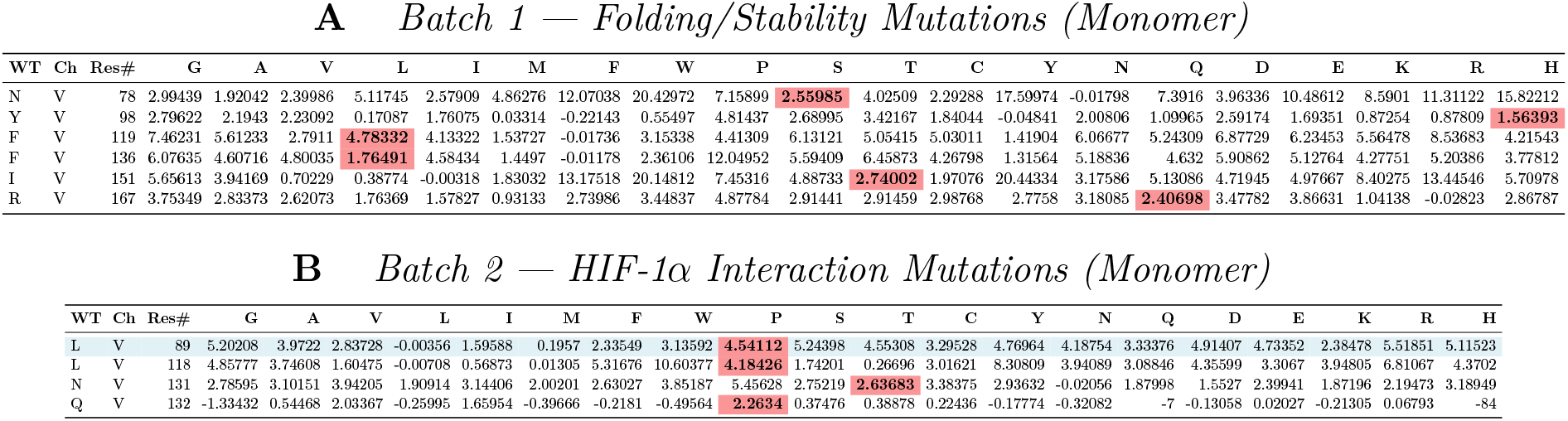
Comparison of energy values for VHL mutations presented in two tables. Table A, Batch 1 mutations (top table) include variants expected to disrupt protein folding and intrinsic structural stability. Mutations highlighted in red: N78S, Y98H, F119L F136L, I151T, and R167Q, exhibit elevated ΔΔG values, indicating significant destabilization of the VHL monomer. Table B, Batch 2 mutations (bottom table) are located at the pVHL–HIF-1*α* binding interface and are predicted to impair protein–protein interactions. Mutations highlighted in red: L89P, L118P, N131T, and Q132P, show high ΔΔG values, suggesting disruption of binding affinity or structural compatibility. L89P is additionally highlighted in blue to emphasize its particularly strong destabilizing effect at the interaction surface.

Batch 1 mutations (Table 2A) are primarily associated with disruptions in protein folding and intrinsic structural stability. These include N78S, Y98H, F119L, F136L, I151T, and R167Q (highlighted in red in Table 2A). All of these mutations exhibit elevated ΔΔG values, with N78S, F119L, and I151T showing the most substantial increases, indicating severe destabilization of the VHL monomer. The high ΔΔG values of F119L and F136L likely result from disruption of hydrophobic core packing, while Y98H, which shows a moderate ΔΔG increase, aligns with its known role as a cancer-associated hotspot. These findings are consistent with previous studies linking such mutations to impaired folding and accelerated degradation [6, 7].

Batch 2 mutations (Table 2B) are predicted to disrupt the interaction between pVHL and HIF-1*α*. These include L89P, L118P, N131T, and Q132P (highlighted in red in Table 2B). L89P is additionally highlighted in blue to emphasize its particularly strong destabilizing effect at the interaction surface. All four mutations showed elevated ΔΔG values at critical interface residues, suggesting that these changes impair the binding affinity or structural alignment required for stable complex formation. This supports a model in which binding interface mutations lead to functional loss through interaction disruption [6].

Together, the ΔΔG profiles in Table 2 support the functional classification of VHL mutations and offer insight into the distinct molecular mechanisms driving monomer destabilization and interaction failure.

#### 3.2.1 Heatmap Representation of VHL Monomer Results

To visualize the predicted effects of each mutation on VHL monomer stability, a heatmap of FoldX-calculated ΔΔG values was generated (Figure 3). Columns represent individual mutations, while rows denote the substituted amino acids. The color gradient reflects the degree of destabilization, with darker red indicating higher ΔΔG values.

**Figure 3:**
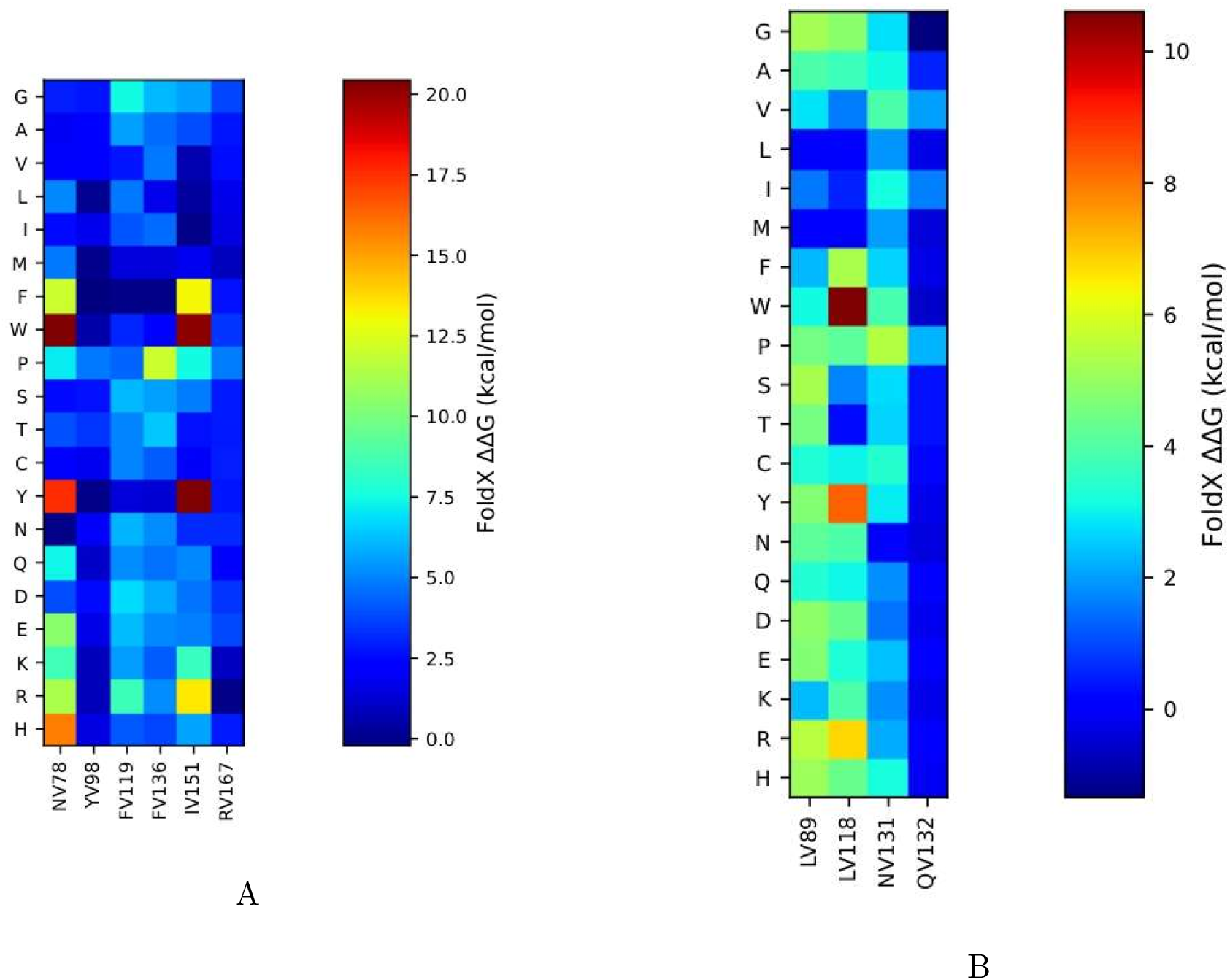
Heatmap comparison of mutation data for the two mutation groups. Figure A on the left shows the heatmap for Batch 1 mutations, which primarily affect protein folding or stability. Figure B on the right displays the heatmap for Batch 2 mutations, which mainly disrupt the interaction between VHL and HIF-1*α*.

Panel A highlights Batch 1 mutations, which show broader and more intense destabilization profiles, consistent with their role in disrupting protein folding and structural integrity. Certain substitutions, such as tryptophan or proline, produced particularly high ΔΔG values across multiple sites. Panel B shows Batch 2 mutations, which mainly affect protein-protein interaction. Although these exhibit milder destabilization overall, a few substitutions still cause significant local effects. This supports the view that the two mutation groups impair VHL function through distinct structural mechanisms.

#### 3.2.2 PyMOL Visualization of VHL Monomer Stability Results

To complement the thermodynamic predictions for the monomer, a structural analysis was conducted using PyMOL to examine the local effects of critical VHL mutations on monomer stability. A representative case here is the L89P mutation, which involves replacing leucine, a hydrophobic, aliphatic residue embedded stably in a *β*-sheet core, with proline, a cyclic amino acid known for its rigidity and its tendency to disrupt regular secondary structures.

In the wild-type conformation (Figure 4A), leucine at position 89 (highlighted in blue) is buried within a hydrophobic pocket and is well accommodated by surrounding side chains (shown in orange), maintaining optimal spatial arrangement without steric interference. This configuration supports the proper folding and structural integrity of the *β*-sheet.

**Figure 4:**
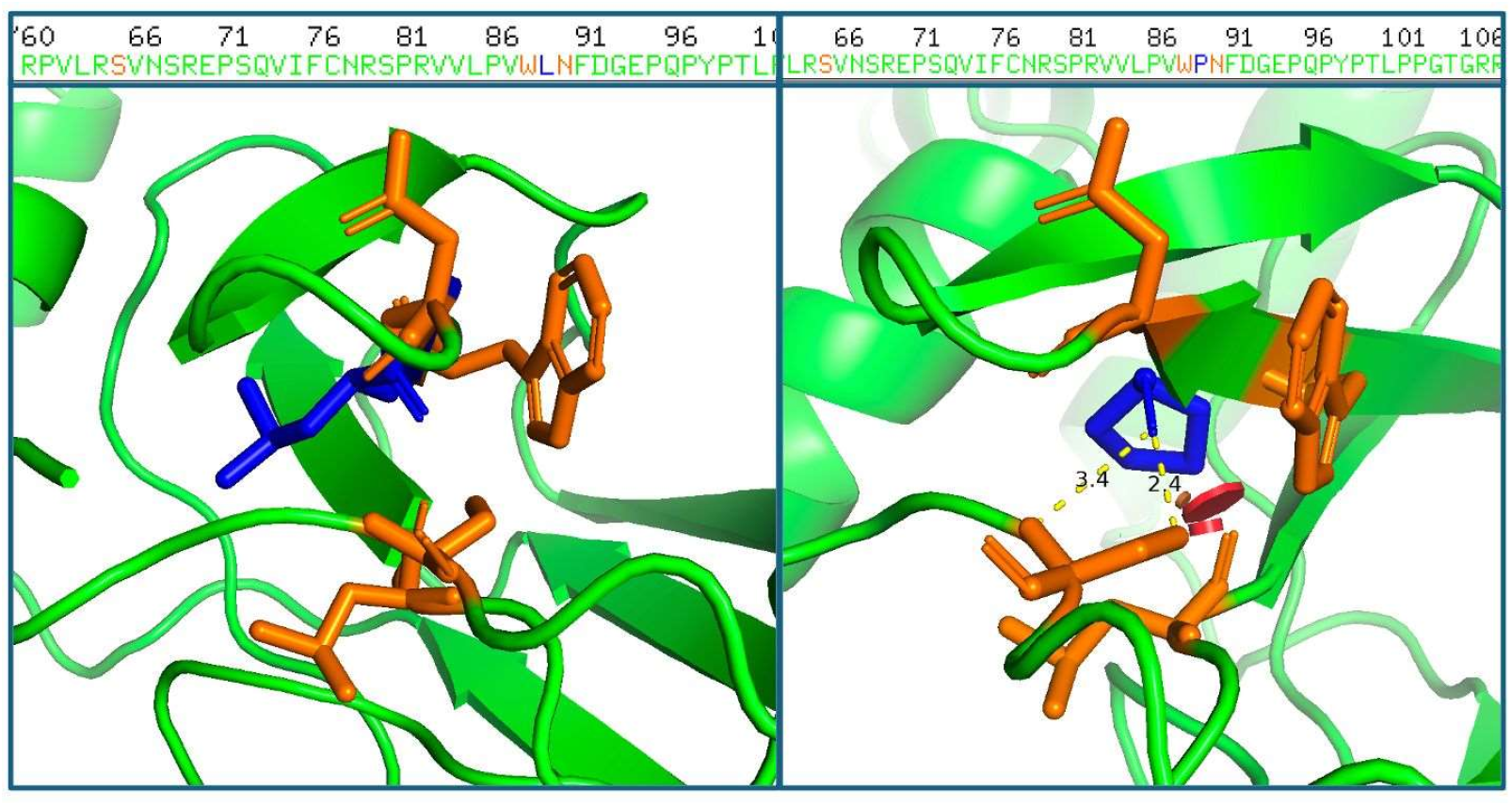
Structural comparison of the VHL protein highlighting the impact of the L89P mutation, visualized using PyMOL. (A) Wild-type structure showing leucine at position 89 (blue), buried within a hydrophobic core formed by surrounding side chains (orange). The residue is stably packed, with no evidence of steric clashes, supporting a well-folded local structure. (B) Mutant structure in which leucine is replaced by proline, a rigid, cyclic residue known to induce kinks and restrict backbone flexibility. The substitution causes repositioning of neighboring hydrophobic side chains (orange), reducing the distance between atoms to below 4 Å and introducing potential steric clashes. This is visualized by a red disk, indicating atomic overlap and conformational strain. The distortion likely destabilizes the *β*-sheet architecture and compromises monomer stability.

In contrast, the Batch 2 mutation (Figure 4B) shows significant local distortion following the introduction of proline. As a result of its rigid ring structure, proline imposes constraints on the backbone dihedral angles, causing a kink that disrupts the native *β*-sheet geometry. This distortion forces adjacent residues into strained conformations, reducing interatomic distances to as close as 2.4–3.4 Å, below the threshold for favorable van der Waals interactions (4 Å). These atomic clashes are highlighted by a red disk (Figure 4B), representing steric hindrance and physical overlap between atoms. The resulting strain destabilizes the local fold and correlates with the elevated ΔΔG values predicted by computational analysis.

Similar structural disruptions were observed for other Batch 2 mutations, such as Q132P and N131T, which are located near the HIF-1*α* binding interface. These mutations are predicted to affect the flexibility and spatial alignment of the backbone, both of which are crucial for effective protein–protein interactions. As a result, these changes could impair VHL complex formation and contribute to the functional loss of VHL [6].

### 3.3 Binding Free Energy and Stability – Of the (VHL-HIF) complex

For the analysis of the complex, the structure was from PDB entry – 4AJY with the solvent and other chains removed, such that only the V (VHL protein) and H (HIF protein) chains remained. Some extra small molecules were also selected and removed.

#### 3.3.1 Simulation in PyMol and Naccess

Visual inspection of the complex in PyMOL (Figure 5), along with distance measurements to the HIF protein, supports the same conclusion for other residues such as N131 and G132. Although these residues are not buried in the core, they are positioned far from the interaction interface, suggesting they are also unlikely to play a direct role in the binding interaction.

**Figure 5:**
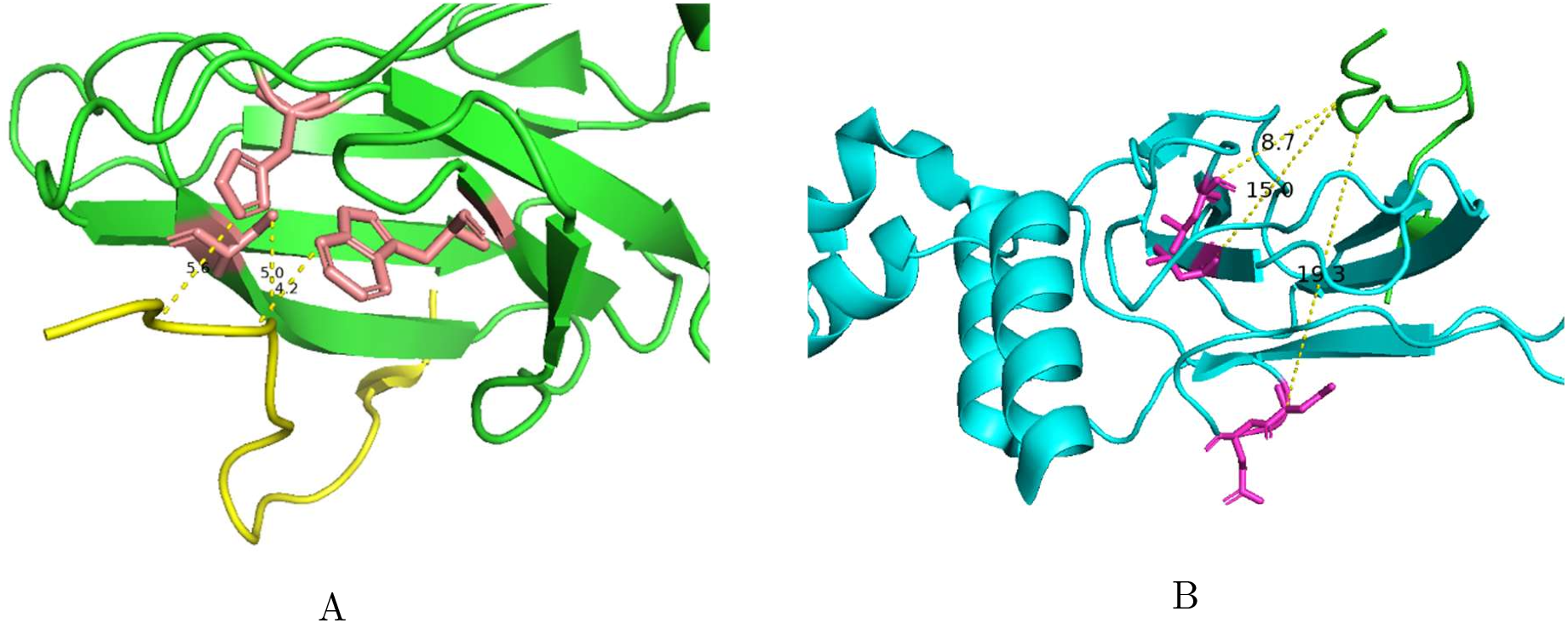
Structural positions of interface-related mutations in a protein–protein complex. A: Mutations (pink) located close to the interface between two proteins (green and yellow), with interatomic distances indicating direct contact. B: Mutations (pink) positioned farther from the interface (cyan and green proteins), with measured distances (e.g., 8.7 Å, 15.0 Å) suggesting indirect effects on binding.

Naccess estimates the solvent-accessible surface area of atoms by simulating the rolling of a spherical probe across the van der Waals surface of a macromolecule. When applied to the monomeric unit, this analysis showed that residues L89 and L118 have side-chain solvent accessibilities of only 0.1% relative to their fully exposed states. This extremely low value indicates that both residues are deeply buried within the hydrophobic core of the protein, making them structurally inaccessible and unlikely to participate in interactions at the protein–protein interface.

#### 3.3.2 Simulating Mutations and Resulting ΔΔG values

Batch 2 mutations—L89P, L118P, N131T, and Q132P—occur further from the interaction interface (Figure 5B). FoldX predicted low ΔΔG values for these substitutions, suggesting minimal impact on global protein stability. These results are also presented in Table 3 – Top Table.

**Table 3:**
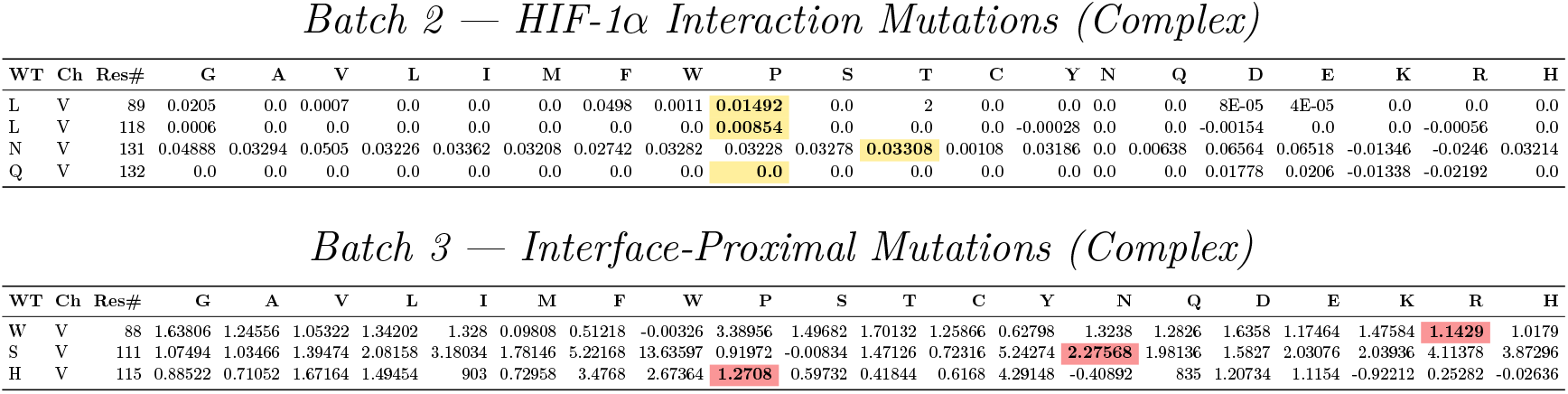
Comparison of energy values for VHL-HIF mutations presented in two tables. Batch 2, top displays the energy values, which are primarily associated with mutations far from the interface. Notably-L89P, L118P, N131T, and Q132P have low ΔΔG values. Batch 3, bottom presents energy values for mutations close to the VHL and HIF-1*α* interface. Destabilizing mutations— W88R, S111N, and H115P - are highlighted in red, reflecting the highest energy values and indicating significant interaction disruption.

In contrast, Batch 3 mutations—W88R, S111N, and H115P—are located near the interface of the VHL complex (Figure 5A). As expected, FoldX predicted high ΔΔG values for these variants, indicating significant destabilization. These values are detailed in Table 3 – Bottom Table and accompanying CSV file and can be visualized in a heatmap like for the monomer (not shown).

It is also worthy to note that other mutations of the same residues could be more destabilizing (eg. W88P, H115W) or even stabilizing (eg. H115N)

### 3.4 Molecular Dynamics and Ensemble Mutational Investigation

#### 3.4.1 Generation of an Ensemble of Conformations for the Wild-Type Structure of the VHL and VHL-HIF Complex and Structural Visualization

To investigate the conformational dynamics of the VHL protein, ten models were generated using the CABSFlex simulation tool. Both the VHL monomer and the VHL-HIF complex were analyzed in this phase of the study. The CABSFlex output files, along with the corresponding Root Mean Square Fluctuation (RMSF) plots, are presented in Figure 6. Additionally, RMSD values were calculated by aligning each model to the reference monomer structure, providing insight into the structural deviations observed across the ensemble.

**Figure 6:**
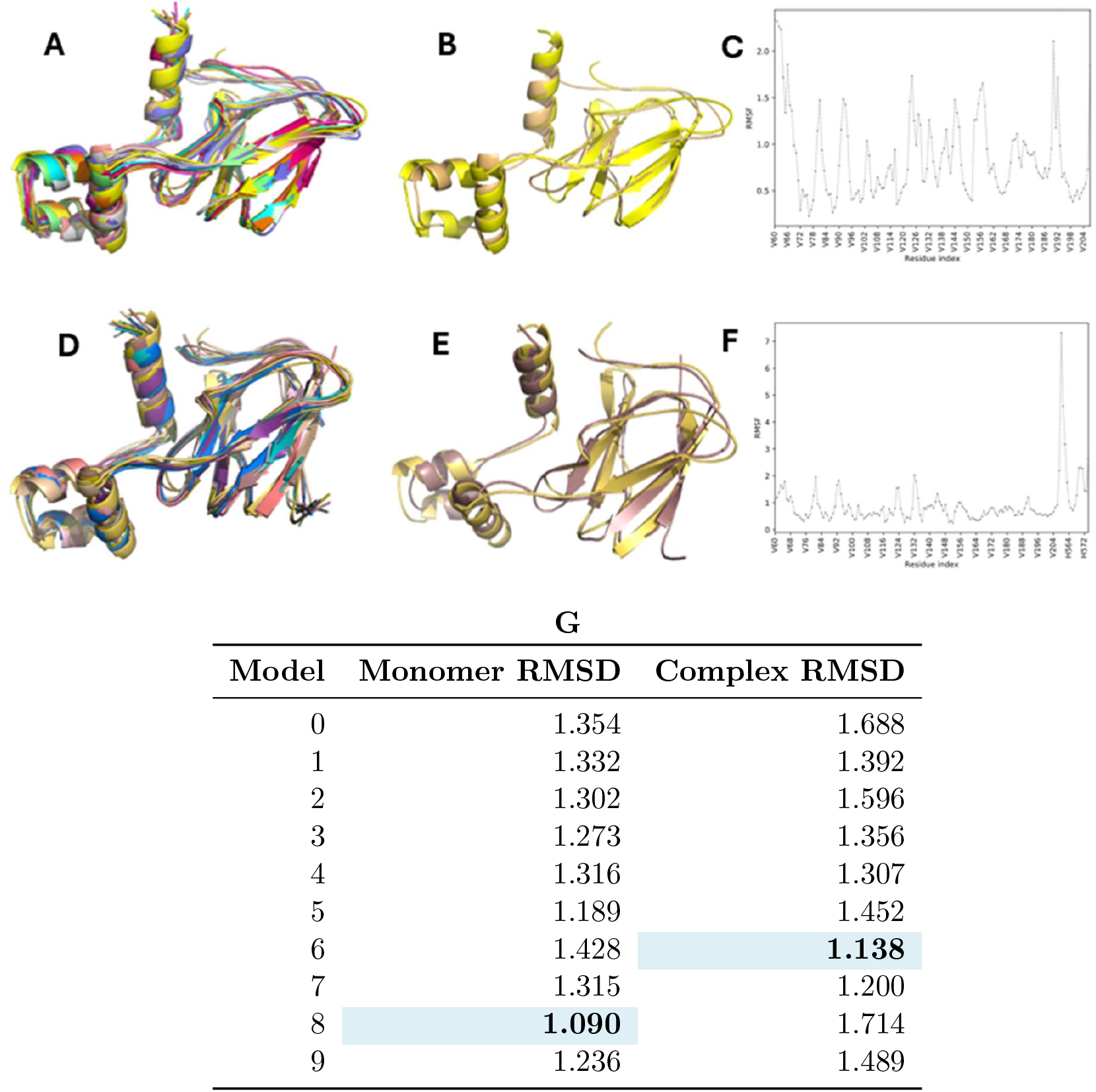
Visual representation of CABSFLEX models and RMSF and RMSD results upon alignment: A) CABSFELX models for VHL monomer aligned to the static structure in PyMol; B) Model 8 of VHL monomer aligned to the static structure in PyMol; C) RMSF plot for the dyna dynamic simulation of VHL monomer; D) CABSFELX models for VHL and HIF complex aligned to the static VHL monomer structure in PyMol; E) Model 6 of VHL and HIF complex aligned to the static monomer structure in PyMol; F) RMSF plot for the dynamic simulation of VHL and HIF complex; G) RMSD values upon alignment of VHL monomer models generated with MC method and VHL and HIF complex models generated with MC method to the VHL static monomer structure.

Figure 6A illustrates the dynamic behavior of the VHL monomer throughout the simulation. While some secondary structural elements are retained across most models, several regions— particularly those composed of *β*-sheets and short turns—experience notable structural degradation. This is especially pronounced in the regions spanning residues 170–177, 92–97, 146–150, 104–111, and 71–78, where short *β*-strands and the N-terminal segments of *α*-helices tend to unfold, indicating localized structural instability. In some cases, these *β*-strands are entirely lost during the simulation. These regions are likely to experience natural folding and unfolding transitions in the protein’s native state, and such dynamic behavior may be intrinsic to VHL’s functional properties. Notably, loop regions show the highest degree of conformational variability across the models, which aligns with their known structural flexibility. This observation is further supported by the RMSF (Root Mean Square Fluctuation) plot, which highlights increased displacement for residues located within these loop regions, underscoring their dynamic nature. Despite this partial loss of secondary structure, the simulation remains informative and biologically relevant. Among the models, Model 8 exhibits the highest accuracy, with a root-mean-square deviation (RMSD) of 1.09.

Figure 6 also shows how the VHL-HIF complex behaves, and it’s clear that it is much more stable than the monomer form. The main secondary structures, especially the *β*-sheets, stay well-formed in almost all of the models. While some loop regions—particularly around residues 93–97 and 103–106—still show some movement, most of the structure stays consistent. This added stability likely comes from the way HIF binds to VHL, especially in the region between residues 91 and 154, which includes the important *β*-sheet area [13].

Just like in the monomer, there are still flexible parts in the complex, but there are fewer flexible secondary structures. The area where HIF binds shows less movement, which suggests that this interaction helps keep the protein folded properly and stable.

Among the models analyzed, Model 6 has the lowest RMSD value (1.138), which means it is the most stable and accurate model in the group.

#### 3.4.2 Mutational Scan Using the CABSFLEX Structures and Comparison with the Experimental Structure

To further explore the dynamics of the VHL monomer, a mutational scan was performed using Model 8, selected based on its low RMSD (1.09 Å), indicating high structural accuracy. This self-scan serves to evaluate the internal stability of the model and to assess its suitability for downstream analyses. Overall, the model appears structurally sound, with the majority of residues showing moderate stability upon mutation. However, residue H125 stands out, exhibiting a notably high ΔΔG of 3.8 kcal/mol upon self-mutation, suggesting potential local instability or a critical structural role. This change in free energy may be an indicator of low accuracy and existence of steric crashes or unfavorable interactions that the local sampling of FoldX cannot resolve. No mutations resulted in a free energy change lower than –1 kcal/mol.

Since Model 8 did not exhibit significant structural inaccuracies, a mutational analysis was conducted on the CABS-flex–generated ensemble. The results of this analysis were aggregated and are presented in Figure 7. Upon mutation of selected residues, several positions exhibited notable changes in free energy, reflecting potential impacts on protein stability.

**Figure 7:**
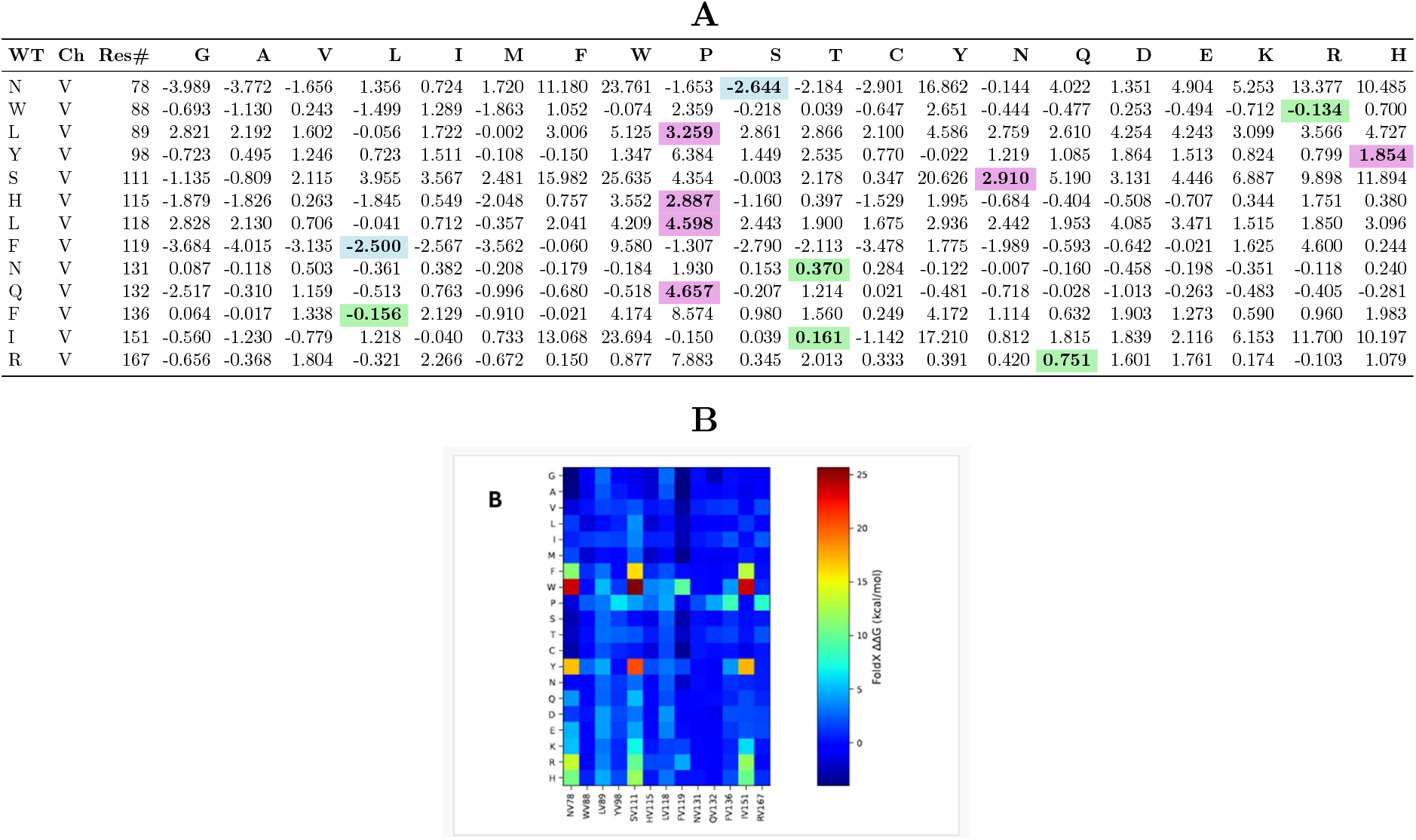
Comparison of energy values for VHL model 8 ensemble mutations with MutateX : A) Numerical values for the average finals of mutational changes with evidentiated free energy change for mutations reported in literature: blue – negative energy values associated with stability of the mutation; green – mutations with values ±1 kcal/mol which doesn’t show any significant change in stability; purple – mutations with positive high energy values that show instability; B) Visual representation of energy changes as a heatmap.

The results in Figure 7 are also consistent with investigations into the static structure (Figure 3) discussed above. In both investigations, residues such as W88, H115, L118, F119, N131, Q132, F136, and R167 generally appeared to be stable, although some non-stabilizing mutations were observed. In contrast, residues including N78, L89, Y98, S111, and I151 demonstrated highly destabilizing responses, with certain mutations - specifically N78W, S111W, and I151W - exceeding +20 kcal/mol in ΔΔG. These results suggest that substitution with bulky amino acids like tryptophan at structurally constrained or functionally sensitive positions may induce significant unfolding or destabilization of the protein.

Further investigation revealed an interesting discrepancy between simulation results and published data. Some mutations showed a high predicted destabilization in simulations – exceeding +20 kcal/mol - yet were not documented in the scientific literature. However, using AlphaMis-sense [14], these same mutations are classified as likely pathogenic, with scores above 0.9, suggesting they may indeed impact protein function significantly, even in the absence of direct experimental validation.

Conversely, some experimentally validated pathogenic mutations, such as N78S and F119L, showed stabilizing effects in the simulation, with ΔΔG values below –1 kcal/mol. Despite appearing stable based on energetic calculations, these mutations are still flagged by AlphaMis-sense as likely pathogenic and were reported in experimental data as well. This highlights that a mutation can still impair protein function despite maintaining or even improving thermodynamic stability. A single amino acid change, especially in a functionally critical region, can disrupt interactions, binding affinity, or regulatory roles without necessarily destabilizing the protein’s overall fold.

## 4 Discussion

Loss-of-function mutations in the VHL tumor suppressor gene lead to the accumulation of HIF-1*α*, promoting uncontrolled cell growth and tumorigenesis. VHL normally binds to hydroxylated HIF-1*α*, targeting it for degradation and preventing its activation of proliferative and angiogenic genes.

VHL mutations lead to a predisposition to tumors and can therefore lead to cancer. Here, we analysed the mutations previously reported in the literature and predicted to be likely pathogenic by AlphaMissense.

### 4.1 Widespread Pathogenicity Indicates Structural and Functional Importance of the Central VHL Region

Not only the selected mutations, but also many residues within the 60 to 200 region of the VHL protein show high AlphaMissense pathogenicity scores. This region includes the *β*-domain and the HIF-binding domain, which are essential for the tumor suppressor function of VHL. The frequency of likely pathogenic variants in this region indicates their important role in the function of the VHL protein.

### 4.2 Structural Stability and Simulation of the VHL Monomer

The classification of VHL monomer missense mutations into two functional categories, those affecting intrinsic structural stability (Batch 1) and those impairing protein–protein interactions (Batch 2), provides a mechanistic framework for understanding their pathogenic consequences. Batch 1 mutations (N78S, Y98H, F119L, F136L, I151T, R167Q) primarily compromise protein folding and core stability, whereas Batch 2 mutations (L89P, L118P, N131T, Q132P) target the interface between pVHL and its key substrate, HIF-1*α*, which is critical for hypoxia signaling and tumor suppression.

FoldX-based ΔΔG calculations revealed significant destabilization induced by several Batch 1 mutations, particularly N78S and I151T, which aligns with their association with misfolding and degradation in prior studies [6, 7]. These results highlight the vulnerability of the protein’s hydrophobic core and *β*-sheet structure to changes in residue size or chemical properties. Y98H, a recognized cancer hotspot, displayed a moderate destabilizing effect, suggesting a partial loss of function, consistent with its role in tumorigenesis.

In contrast, Batch 2 mutations, though localized at the protein–protein interface, also yielded elevated ΔΔG values, indicating structural instability at critical interaction sites. L89P and Q132P cause significant local strain, which likely disrupts the geometric and energetic requirements necessary for stable HIF-1*α* binding. Structural analysis via PyMOL supported these findings. For example, the L89P mutation substitutes a buried leucine with a sterically restrictive proline, causing steric clashes and distorting the local *β*-sheet. This distortion reduced interatomic distances to below the van der Waals interaction threshold (~4 Å), a sign of local misfolding and potential failure of interaction with HIF-1*α*.

Similar structural disruptions were observed for other mutations near or at the binding interface, such as Q132P and N131T. These mutations appear to interfere with backbone flexibility and side-chain alignment, further compromising the binding affinity required for proper complex formation. Together, the thermodynamic and structural data indicate that both global misfolding and localized interface disruption contribute to VHL loss-of-function phenotypes.

Amino acid property analysis provided further insight into how changes in physicochemical properties, such as hydrophobicity, rigidity, and polarity, correlate with observed ΔΔG values. The heatmap visualization supported these findings by highlighting mutation-specific energetic trends. Proline substitutions, particularly in Batch 2, were consistently associated with high destabilization, emphasizing the role of rigidity and bulkiness in promoting structural instability.

In conclusion, this integrated computational and structural analysis underscores the critical importance of maintaining both folding stability and precise intermolecular interactions for pVHL functionality. Mutations that impair either aspect likely compromise VHL’s tumor-suppressor role by either promoting degradation or disrupting substrate recognition. These insights not only clarify the molecular basis of VHL-associated diseases but also highlight potential targets for therapeutic intervention through structure-guided drug design or stabilization strategies.

### 4.3 Binding Free Energy and Stability – on the Complex

The high ΔΔG values observed for Batch 3 mutations are consistent with their spatial location near the protein-protein interface, which is particularly susceptible to structural disruption. As illustrated in Figure 5A, W88R introduces a large, charged residue in place of a hydrophobic tryptophan, disrupting local packing. S111N may interfere with hydrogen bonding networks due to its larger polar side chain, and H115P introduces conformational strain, likely distorting nearby secondary structure elements. These effects explain the destabilizing ΔΔG values seen in Table 3, supporting their known roles in disrupting VHL complex stability.

Conversely, Batch 2 mutations, while showing low ΔΔG values, are experimentally known to impair VHL-HIF binding. This discrepancy suggests that these mutations impact function through mechanisms not captured by global stability metrics. For example, L89P, L118P, and Q132P may introduce local rigidity or distort key motifs necessary for HIF recognition, while N131T may subtly affect interaction surfaces. As shown in Figure 5B, these mutations are located away from the interface and do not appear to disrupt core stability, which is reflected in the lower ΔΔG values in Table 3-Top.

So, mutations located near the complex interface showed high ΔΔG values and were predicted by FoldX to be destabilizing, consistent with expectations. In contrast, mutations further from the interface yielded low ΔΔG values, despite being experimentally known to disrupt VHL-HIF binding, highlighting a limitation of stability-based predictions in capturing functionally relevant but structurally subtle effects.

### 4.4 Molecular Dynamics and Ensemble Mutational Analysis

Through molecular dynamics simulations using CABSFlex, the conformational flexibility of both the VHL monomer and the VHL-HIF complex was explored. The VHL monomer showed notable instability in several *β*-sheet and loop regions, particularly between residues 71–177, reflecting natural flexibility likely linked to its functional role. In contrast, the VHL-HIF complex displayed greater structural stability, especially within the HIF-binding domain (residues 91–154), suggesting that interaction with HIF helps maintain VHL’s folded structure.

Among the generated models, Model 8 (monomer) and Model 6 (complex) showed the lowest RMSD values, indicating the highest structural accuracy. Model 8 was used for mutational analysis, revealing that while most residues were stable, mutations at certain positions like H125 led to significant energy increases, indicating potential structural strain.

A broader mutational scan identified several residues (e.g., N78, S111, I151) where substitutions, especially with bulky residues like tryptophan, caused major destabilization (*>*+20 kcal/mol). Interestingly, discrepancies emerged when comparing simulation results with data from AlphaMissense [14] and experimental literature. Some mutations predicted as stable in simulations (e.g., N78S, F119L) were still labeled as likely pathogenic in AlphaMissense and validated experimentally. This suggests that pathogenicity can result from subtle functional disruptions, not just loss of structural stability.

Together, these findings emphasize the value of combining structural simulations with machine learning predictions and experimental data to better understand the impact of mutations on protein function.

## 5 Conclusions

This project provides an in-depth analysis of VHL disease, demonstrating how computational methods can help reveal its genetic drivers and functional implications. Molecular dynamics simulations and mutational analyses offered important insights into the structure and behavior of the VHL protein. While the VHL monomer demonstrated considerable structural flexibility, its interaction with HIF resulted in more stability. Mutational scanning identified both stabilizing and destabilizing variants, although some mutations showed discrepancies between predicted structural stability and known pathogenic outcomes. These results suggest that structural stability alone is not fully responsible for the pathogenicity of mutations, emphasizing the importance of integrative approaches in assessing protein function and disease relevance. Overall, these data-driven studies show how mutations in the VHL tumor suppressor gene contribute to tumor development and progression. Bioinformatics tools not only deepen our molecular understanding of disease mechanisms but also provide the basis for new diagnostic and therapeutic strategies.

## Acknowledgements

We thank Elena Papaleo (Technical University of Denmark) for her guidance and supervision throughout this project.

## References

[1] J. R. Forman, C. L. Worth, G. R. J. Bickerton, T. G. Eisen and T. L. Blundell, “Structural bioinformatics mutation analysis reveals genotype-phenotype correlations in von Hippel-Lindau disease and suggests molecular mechanisms of tumorigenesis,” Proteins, vol. 77, no. 1, p. 84–96, 2009.

[2] G. L. Semenza, “Hydroxylation of HIF-1: oxygen sensing at the molecular level,” Physiology, vol. 19, pp. 176–182, 2004.

[3] K. Degn, L. Beltrame, M. Tiberti and E. Papaleo, “PDBminer to Find and Annotate Protein Structures for Computational Analysis,” Journal of chemical information and modeling, vol. 63, no. 23, p. 7274–7281, 2023.

[4] F. Chesnel, A. Couturier, A. Alusse, J.-P. Gagné, G. G. Poirier and D. Jean, “The prefoldin complex stabilizes the von Hippel-Lindau protein against aggregation and degradation,” PLoS genetics, vol. 16, no. 11, 2020.

[5] M. D. Shmueli, L. Schnaider, D. Rosenblum, G. Herzog, E. Gazit and D. Segal, “Structural insights into the folding defects of oncogenic pVHL lead to correction of its function in vitro,” PloS one, vol. 8, no. 6, 2013.

[6] C. Core, “VHL N78S - Gene Variant Detail,” [Online]. Available: https://ckb.genomenon.com/geneVariant/show?geneVariantId=9008&tabType=EXTENDED_EVIDENCE&utm_source=chatgpt.com.

[7] S. Buart, S. Terry, M. K. Diop, P. Dessen, S. Couvé, A. Abdou, J. Adam, J. Thiery, P. Savagner and S. Chouaib, “The most common VHL point mutation R167Q in hereditary VHL disease interferes with cell plasticity regulation,” Cancers, vol. 13, no. 6, p. 13897, 2021.

[8] K. Steward, “Essential Amino Acids: Chart, Abbreviations and Structure,” 2025. [Online]. Available: https://www.technologynetworks.com/applied-sciences/articles/essential-amino-acids-chart-abbreviations-and-structure-324357.

[9] A. Kuriata, A. M. Gierut, T. Oleniecki, M. P. Ciemny, A. Kolinski, M. Kurcinski and S. Kmiecik, “CABS-flex 2.0: a web server for fast simulations of flexibility of protein structures,” Nucleic Acids Research, vol. 46, no. W1, p. W338–W343, 2018.

[10] Schrodinger, LLC. 2010. The PyMOL Molecular Graphics System, Version 1.3.

[11] M. Tiberti, T. Terkelsen, K. Degn, L. Beltrame, T. C. Cremers, I. d. Piedade, M. D. Marco, E. Maiani and E. Papaleo, “MutateX: an automated pipeline for in silico saturation mutagenesis of protein structures and structural ensembles,” Briefings in Bioinformatics, vol. 23, no. 3, 2022.

[12] J. Schymkowitz, J. Borg, F. Stricher, R. Nys, F. Rousseau and L. Serrano, “The FoldX web server: an online force field,” Nucleic Acids Res., vol. 33, 2005.

[13] K. Tanimoto, Y. Makino, T. Pereira and L. Poellinger, “Mechanism of regulation of the hypoxia-inducible factor-1 alpha by the von Hippel-Lindau tumor suppressor protein,” EMBO J., pp. 4298–4309, 2000.

[14] J. Cheng, “Accurate proteome-wide missense variant effect prediction with AlphaMissense,” Science, vol. 381, 2023.

